# Structures of MmpL complexes reveal the assembly and mechanism of this family of transporters

**DOI:** 10.1101/2025.06.22.660944

**Authors:** Zhemin Zhang, Rakesh Maharjan, William D. Gregor, Philip A. Klenotic, Edward W. Yu

## Abstract

We co-expressed the MmpL5 transporter and MmpS5 adaptor proteins in *Mycobacterium smegmatis* and defined their structures from these detergent-solubilized crude membranes. Data generated from these samples allowed us to simultaneously solve three distinct classes of membrane protein complexes to high resolutions. We observed that MmpL5 presents as a monomer in complex with the cytosolic meromycolate extension acyl carrier protein M (AcpM) in a molar ratio of 1:1, where these AcpM-MmpL5 complexes closely pack together to generate regular two-dimensional arrays. We also identified MmpL5 as a trimer that interacts with MmpS5 and AcpM in a molar ratio of 3:3:3 to assemble the tripartite complex AcpM-MmpL5-MmpS5 that spans both the inner and outer membranes of the mycobacterium. In addition, we discovered that MmpL5 and AcpM are able to form the trimeric AcpM-MmpL5 complex in a molar ratio of 3:3. The structural data reveal that the full-length MmpL5 trimer is capable of spanning the entire mycobacterial cell envelope to transport substrates. However, this assembly requires the presence of MmpS5 to stabilize secondary structural features of the MmpL5 periplasmic subdomains.

## Introduction

Tuberculosis (TB), an airborne infectious disease, is caused by the causative agent *Mycobacterium tuberculosis* (Mtb). It has returned to being the leading cause of death from a single infectious agent, where this position was briefly defeated by the coronavirus during the COVID-19 pandemic. In 2023, it was estimated that 10.8 million people fell ill with TB and approximately 1.25 million patients died from this disease worldwide (*1*). TB is currently the leading cause death (*1*), exceeding both malaria and HIV/AIDS, causing almost twice as many deaths as the latter (*2*). To date, approximately 1/3 of the world’s population is infected by Mtb with most having the latent form of disease; however, approximately 10% of this population will progress to active TB (*3*).

The cell envelope of Mtb appears to be one of the most complex membranes of all bacteria and plays a dominant role in Mtb’s pathogenesis and virulence. The outer mycomembrane is very rigid and extremely impermeable. It provides a waxy, hydrophobic layer and contributes to creating a strong barrier against many antimicrobials as well as the host’s immune response (*4*). The inner leaflet of the outer mycomembrane mainly constitutes very long chain (C60-C90) α-alkyl β-hydroxy fatty acids named mycolic acids (MAs), which are either covalently linked to the arabinogalactan-peptidoglycan layer as mycolyl arabinogalactan peptidoglycans (mAGPs) or incorporated into trehalose dimycolates (TDMs). The outer leaflet of the mycomembrane contains several non-covalently associated lipids, such as phthiocerol dimycocerosates (PDIMs), sulfolipids (SLs) and trehalose monomycolates (TMMs) (*4*).

The mycobacterial membrane protein large (MmpL) transporters (*5*), belonging to a subfamily of the resistance-nodulation-division (RND) superfamily of membrane proteins (*6*), are major contributors to mycobacterial cell wall biogenesis. The genome of Mtb encodes 13 MmpL transporters (*5*), with many of them engaged in the export of fatty acids and lipid components to the mycobacterial cell envelope. For example, MmpL3 is essential for the translocation of TMM to the mycobacterial surface (*7–9*), whereas MmpL11 is responsible for delivering very-long chain triacylglycerols (LC-TAG) and mycolate wax ester (MWE) (*10, 11*). MmpL7 and MmpL8 are involved in the transport of virulence-associated lipids PDIM and SL-1 (*12, 13*), and MmpL10 contributes to shuttle diacyltrehalose to the cell envelope (*14*). MmpL4 and MmpL5 share redundant functions in the export of siderophore lipids, such as mycobactins and carboxymycobactins, and are required for iron acquisition (*15*).

To elucidate the molecular mechanisms of these MmpL membrane proteins, we and others previously solved high-resolution structures of the MmpL3 transporter (*16–19*). We also determined the structure of MmpL3 bound with the lipid TMM, revealing two substrate binding sites that facilitate the transport of TMM, or other lipids, to the periplasm (*20*). In addition, we resolved structures of both MmpL4 and MmpL5, which employ similar transport systems to export siderophores for iron scavenging (*21*), as well as the structure of MmpL5 bound with meromycolate extension acyl carrier protein M (AcpM), indicating that MmpL5 is capable of specifically binding AcpM to form the stable AcpM-MmpL5 complex (*22*).

Although these MmpL3, MmpL4 and MmpL5 structures all present monomeric architectures of the transporters, their oligomerization states and assemblies are still under debate. A homotrimeric organization of both the MmpL3 and MmpL5 transporters has been proposed based on negative staining and single-molecule imaging (*14, 23*). Additionally, MmpL transporters have been demonstrated to interact with various proteins *in vivo* (*24, 25*), in particular their corresponding MmpS adaptor membrane proteins. However, the roles of these accessory proteins in relation to substrate transport remain unknown. It is not known how these MmpL transporters form complexes with their MmpS adaptors. It is also unclear if these MmpL transporters by themselves are sufficient for the shuttling of substrates from the inner membrane to the periplasm and/or across the cell envelope without an intermediate step that passes these substrates to their corresponding interacting partner proteins, although MmpL3 has been implicated to contact the periplasmic α/β-barrel lipoprotein LpqN (*10*) to shuttle TMM, and MmpL5 has been proposed to coordinate with the periplasmic protein Rv0455c (*26, 27*) for siderophore recycling. It is quite obvious that many questions remain with respect to the roles, assemblies, architectures, interactions and molecular mechanisms of the MmpL-MmpS transporter-adaptor systems.

Here we express both the MmpL5 transporter and MmpS5 adaptor in *M. smegmatis* membranes and simultaneously solve high-resolution structures of three different protein complexes from these enriched crude *M. smegmatis* membranes. From this sample, we identified and solved structures of three different MmpL5 membrane protein complexes: the dipartite AcpM-MmpL5 monomeric complex, the tripartite AcpM-MmpL5-MmpS5 trimeric complex and the dipartite AcpM-MmpL5 trimeric complex. Our work provides the first structural insights into the oligomerization, assembly and mechanism of these critical MmpL-MmpS transporter-adaptor systems in mycobacteria.

## Results

We cloned the *M. smegmatis* MmpS5 (MSMEG_1381) and MmpL5 (MSMEG_1382) genes into the pBUN250 expression vector to co-express both membrane proteins in *M. smegmatis* MC^2^-155 cells. However, co-purification of MmpL5 and MmpS5 was unsuccessful, which led us to attempt to obtain structural information of the MmpL5-MmpS5 complex from crude membranes. We collected the *M. smegmatis* MC^2^-155 cell membranes that selectively expressed MmpL5 and MmpS5 and then solubilized them in detergent micelles. The membrane protein components were enriched using sucrose cushion centrifugation. This enriched sample was applied to a holey carbon grid and single-particle cryo-EM images were then collected. Subsequent cryo-EM data were processed using the “Build and Retrieve” (BaR) methodology (*28*). 2D classification of the single-particle images indicated that there were at least three distinct, highly populated classes of images coexisting in this membrane sample. Several iterative rounds of 2D classification allowed us to sort the images into these three classes. Finally, we were able to identify and solve cryo-EM structures of three different membrane protein complexes, corresponding to the dipartite AcpM-MmpL5 monomeric complex, the tripartite AcpM-MmpL5-MmpS5 trimeric complex and the dipartite AcpM-MmpL5 trimeric complex (Fig S1 and Table S1). Mass spectrometry also confirmed the presence of the MmpL5, MmpS5 and AcpM proteins in the enriched sample (Table S2).

### Duplicated AcpM-MmpL5 units cluster together to form two-dimensional arrays

The most abundant single-particle protein molecule found in our sample is a membrane protein complex, where duplications of these complex molecules cluster together and assemble as well-ordered two-dimensional (2D) arrays. A total of 1,037,284 projections were obtained in this protein class. We identified that the arrays are formed by individual, identical repeating units of AcpM-MmpL5, a membrane protein complex consisting of the AcpM meromycolate extension acyl carrier protein and the MmpL5 transporter. We then solved the structure of this complex to a resolution of 3.35 Å (Fig 1A-C, Fig S1 and Table S1).

**Fig 1.**
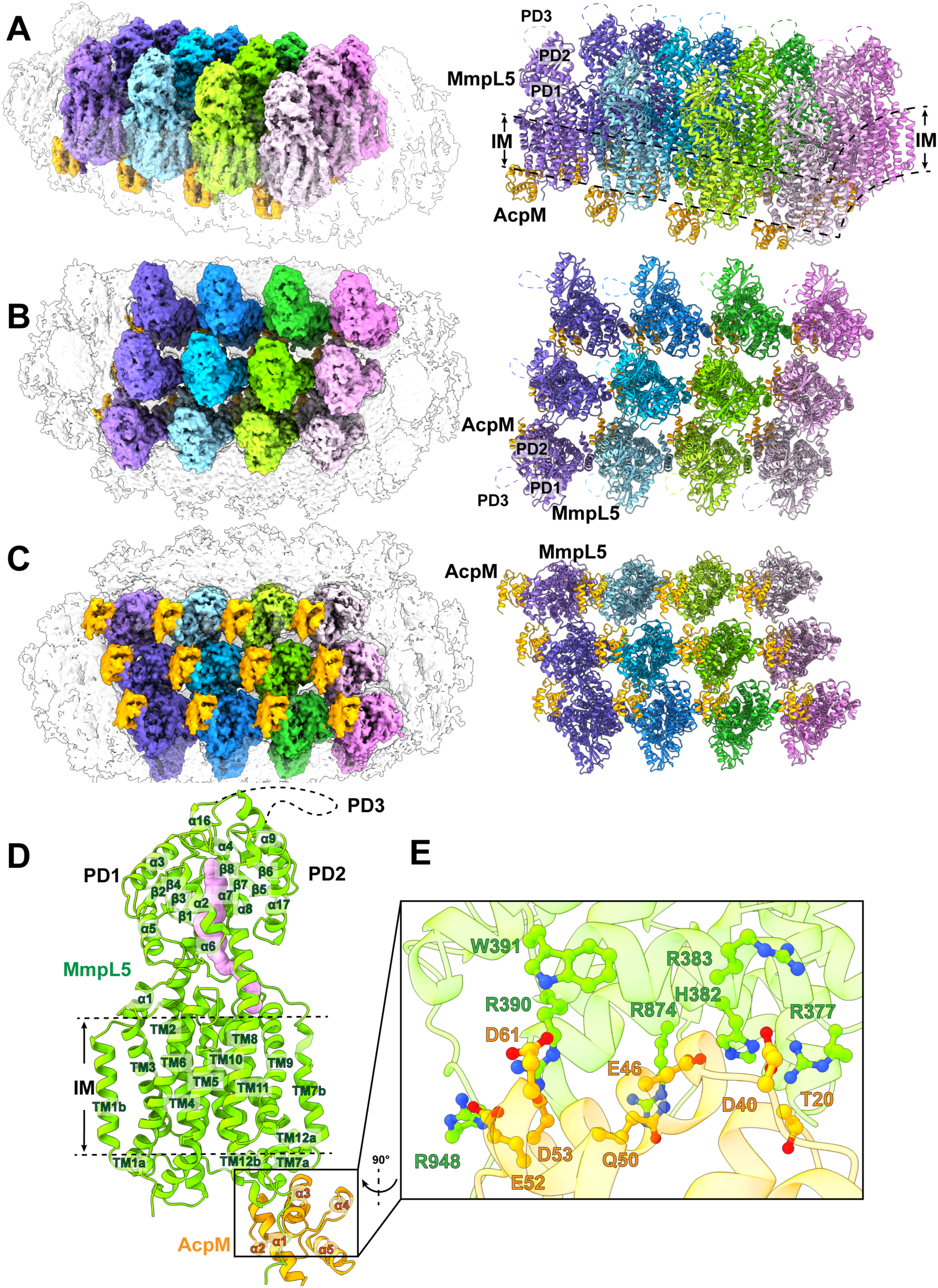
Duplicated AcpM-MmpL5 units form 2D arrays. (A) Side view, (B) top view and (C) bottom view of the cryo-EM maps and ribbon diagrams of duplicated AcpM-MmpL5 units within the 2D arrays. Individual MmpL5 molecules are in different colors. Each AcpM molecule is colored orange. (D) Structure of an individual AcpM-MmpL5 unit within the AcpM-MmpL5 arrays.

The cryo-EM structure indicates that each MmpL5 molecule within the AcpM-MmpL5 arrays is monomeric in oligomerization. Each AcpM-MmpL5 unit is made up of one molecule of MmpL5 and one molecule of AcpM in a molar ratio of 1:1. The structure of each AcpM-MmpL5 unit is nearly identical to our previously published structure of purified AcpM-MmpL5 (*22*). Superimposition of these two structures gives rise to an r.m.s.d. of 0.9 Å (for 725 Cα atoms), suggesting that the conformational states of these two structures are almost the same.

#### Structure of the individual AcpM-MmpL5 unit

MmpL5 contains a transmembrane domain consisting of 12 transmembrane helices (TM1-TM12) and a periplasmic domain composed of three subdomains (PD1-PD3). Within the 2D protein arrays, each MmpL5 molecule indeed constitutes a large transmembrane domain and a large periplasmic domain. Like the purified AcpM-MmpL5 structure determined previously, TM1-TM12 of the transmembrane domain and PD1-PD2 of the periplasmic domain of each MmpL5 transporter within the arrays are clearly defined from the cryo-EM map. However, cryo-EM densities for most residues (residues 493-702) of the periplasmic subdomain PD3 of each MmpL5 molecule are still missing (Fig 1D).

Similar to the purified AcpM-MmpL5 complex structure, each MmpL5 within the 2D arrays forms an elongated channel that spans the pocket surrounded by TMs 7-10 at the outer leaflet of the cytoplasmic membrane and the cavity generated between subdomains PD1 and PD2 in the periplasm (Fig 1D). This elongated channel is presumed to form a path for MmpL5 to transport substrates.

Within each AcpM-MmpL5 repeating unit, AcpM presents a four α-helical bundle structure with a hydrophobic internal core (Fig 1D). Its structure is nearly identical to the previous structure obtained from the purified AcpM-MmpL5 complex (*22*), where superimposition of these two AcpM structures results in an r.m.s.d. of 0.5 Å (for 82 Cα atoms).

#### AcpM-MmpL5 interactions

As with the structure of purified AcpM-MmpL5 (*22*), AcpM interacts predominantly with MmpL5 through charge-charge interactions. The amino acid E46 of AcpM forms a salt bridge with H382 of MmpL5, while E52 and D53 of AcpM create two separate salt bridges with R948 of MmpL5. In addition, residues D40 and D61 of AcpM form two hydrogen bonds with R383 and R390 of MmpL5, respectively. Furthermore, T20, Q50 and D61 of AcpM contact R377, R875 and W391 of MmpL5 via electrostatic interactions to help stabilize this complex (Fig 1E).

### The tripartite AcpM-MmpL5-MmpS5 complex forms a needle-like apparatus that may span the entire mycobacterial cell envelope

The second most abundant protein observed in our cryo-EM grid is an elongated membrane protein complex. We collected a total of 72,455 single-particle projections for this class of images. We identified this membrane protein complex as the tripartite AcpM-MmpL5-MmpS5 complex. The first structural information of this trimeric membrane protein complex was determined to a resolution of 2.81 Å (Fig 2A-D, Fig S1 and Table S1).

**Fig 2.**
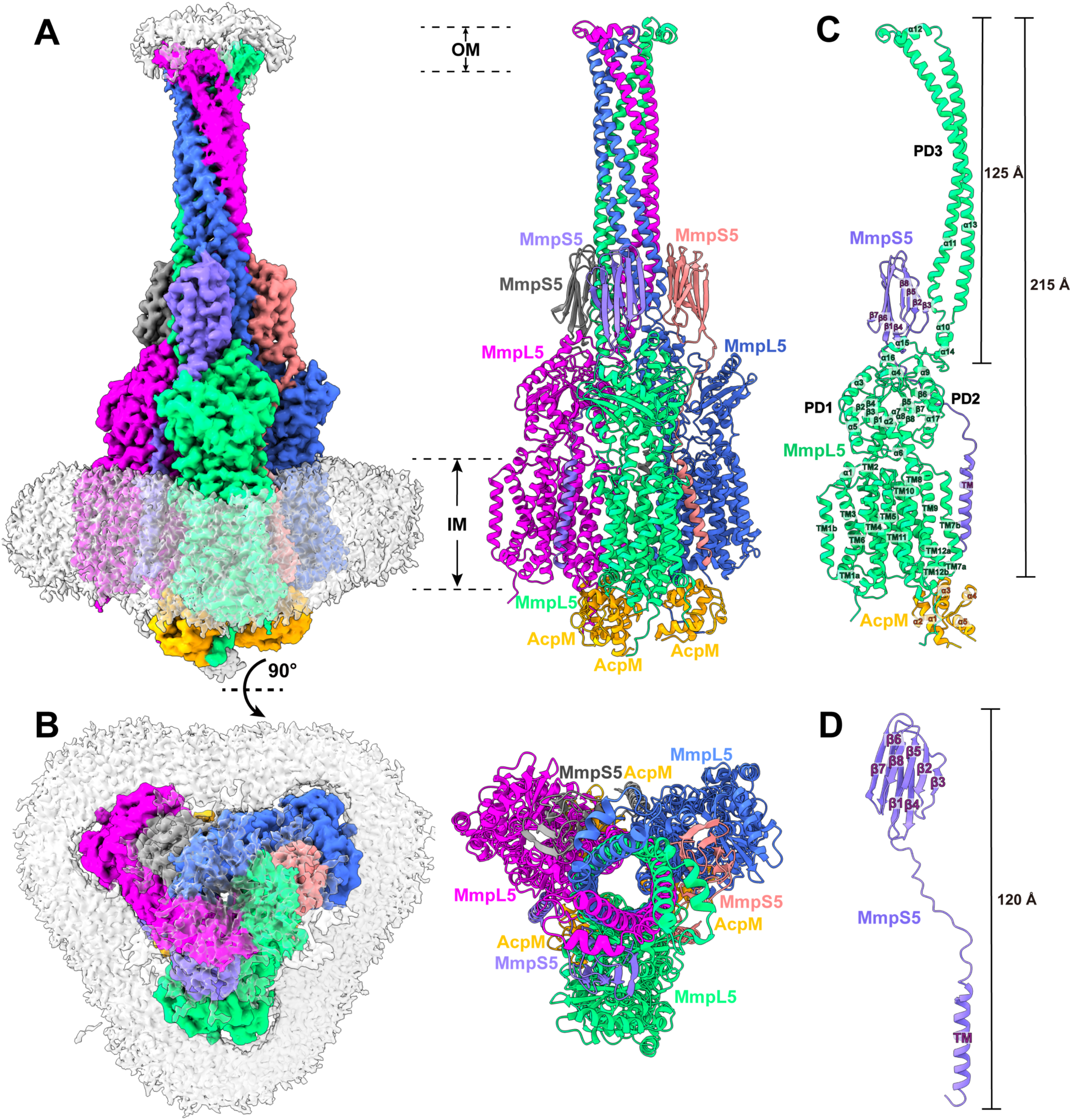
Structure of the tripartite AcpM-MmpL5-MmpS5 complex. (A) Side view and (B) top view of the cryo-EM maps and ribbon diagrams of the AcpM-MmpL5-MmpS5 complex. The structure indicates that MmpL5 assembles as a trimer. The stoichiometry of AcpM:MmpL5:MmpS5 within the complex is in the form of a 3:3:3 molar ratio. Individual MmpL5 and MmpS5 molecules are colored differently. The AcpM molecules are colored orange and the detergent belts in the cryo-EM maps are colored gray. The inner and outer membrane regions are indicated as IM and OM, respectively. (C) Structure of a single unit of AcpM-MmpL5-MmpS5 within the trimeric complex. The structure indicates that the PD3 subdomain of MmpL5 forms an elongated hairpin of 125 Å in length, resulting in a vertical dimension of the MmpL5 protomer to be 215 Å. (D) Structure of a molecule of the MmpS5 adaptor. The structure indicates that MmpS5 is an elongated protein 120 Å long that is composed of an N-terminal transmembrane domain of one single helix (TM) and a C-terminal periplasmic domain of eight β strands (β1-β8).

The structure of the AcpM-MmpL5-MmpS5 complex provides very important information that allows us to significantly fill in our knowledge gap of these MmpL transporters. First, the cryo-EM structure reveals that MmpL5 itself assembles as a trimer within the AcpM-MmpL5-MmpS5 tripartite complex. Second, within the MmpL5 trimer, each MmpL5 protomer uses both its transmembrane and periplasmic domains to contact an MmpS5 adaptor molecule. In addition, each MmpL5 protomer also utilizes its cytoplasmic side to anchor the AcpM acyl carrier protein molecule. Based on the structural information, the stoichiometry of AcpM:MmpL5:MmpS5 within the trimeric complex is in the form of a 3:3:3 molar ratio. Third, subdomain PD3 of each MmpL5 creates an elongated all α-helical hairpin, where the three hairpins specifically contact each other to generate a needle-like, trimeric channel within the MmpL5 trimer. The tip of this trimeric channel is surrounded with detergent molecules, indicating that this tip is embedded in the outer mycomembrane. Therefore, the AcpM-MmpL5-MmpS5 tripartite complex is presumed to span both the inner and outer membrane of the mycobacterium. Fourth, the structure of AcpM-MmpL5-MmpS5 allows us to reveal the first structural information of the full-length MmpS family of adaptor proteins.

#### Structure of the trimeric MmpL5 transporter

Within the AcpM-MmpL5-MmpS5 complex, MmpL5 exists as a homotrimer, where the conformational state of each MmpL5 protomer is identical to each other (Fig 2A-B). Each subunit of MmpL5 is composed of 12 transmembrane helices (TM1–TM12) and a periplasmic domain formed by two periplasmic loops, one between TM1 and TM2 and another between TM7 and TM8. Our cryo-EM density clearly indicates that these two periplasmic loops create three periplasmic subdomains, PD1, PD2 and PD3, of the MmpL5 transporter in this tripartite complex (Fig 2C).

The structural information of TM1-TM12, PD1 and PD2 of each subunit of MmpL5 within the trimer is comparable to that of each individual molecule of MmpL5 residing in the AcpM-MmpL5 2D arrays. Superimposition of the structures of these two MmpL5 protomers, which include the 12 TMs, PD1 and PD2, gives rise to an r.m.s.d. of 1.1 Å, suggesting that the conformations of these two structures are very similar to each other. In addition, the interactions between AcpM and MmpL5 in this tripartite complex are almost identical to those found in the AcpM-MmpL5 2D arrays, where these interactions are dominated by hydrogen-bond, salt-bridge and electrostatic interactions.

#### Structure of the PD3 subdomain

Perhaps, the most exciting structural feature of MmpL5 comes from the PD3 subdomain. Based on the structural information, PD3 is made up of residues 499-684, forming an all α-helical subdomain (α10-α14). A detailed inspection of the structure indicates that this subdomain is mainly created by two elongated helices, α11 and α13, which run antiparallel to each other to form a hairpin. Within the MmpL5 trimer, the three PD3 hairpins intimately contact one another via coiled-coil interactions. The side-by-side packing arrangement of three PD3 subdomains allows a channel-like structure to be created with the center of the channel being formed along the three-fold symmetrical axis (Fig 3A-B). This elongated channel establishes a vertical needle-like shape that forms directly above subdomains PD2 of the three MmpL5 protomers. The needle is approximately 125 Å long and 40 Å wide (Fig. 3A), resulting in a vertical dimension of the MmpL5 trimer to be approximately 215 Å (Fig 2C). This distance is presumed to be long enough to span both the inner and outer membranes of the mycobacterial cell. In addition, residues 579-600, which form the tip of the α-helical hairpin of each PD3 subdomain and the top portion of the trimeric channel, is surrounded with detergent molecules in the cryo-EM map, suggesting that the tip of the trimeric channel is embedded in the membrane layer. Therefore, the assembled MmpL5 trimer itself is able to span the entire mycobacterial cell envelope.

**Fig 3.**
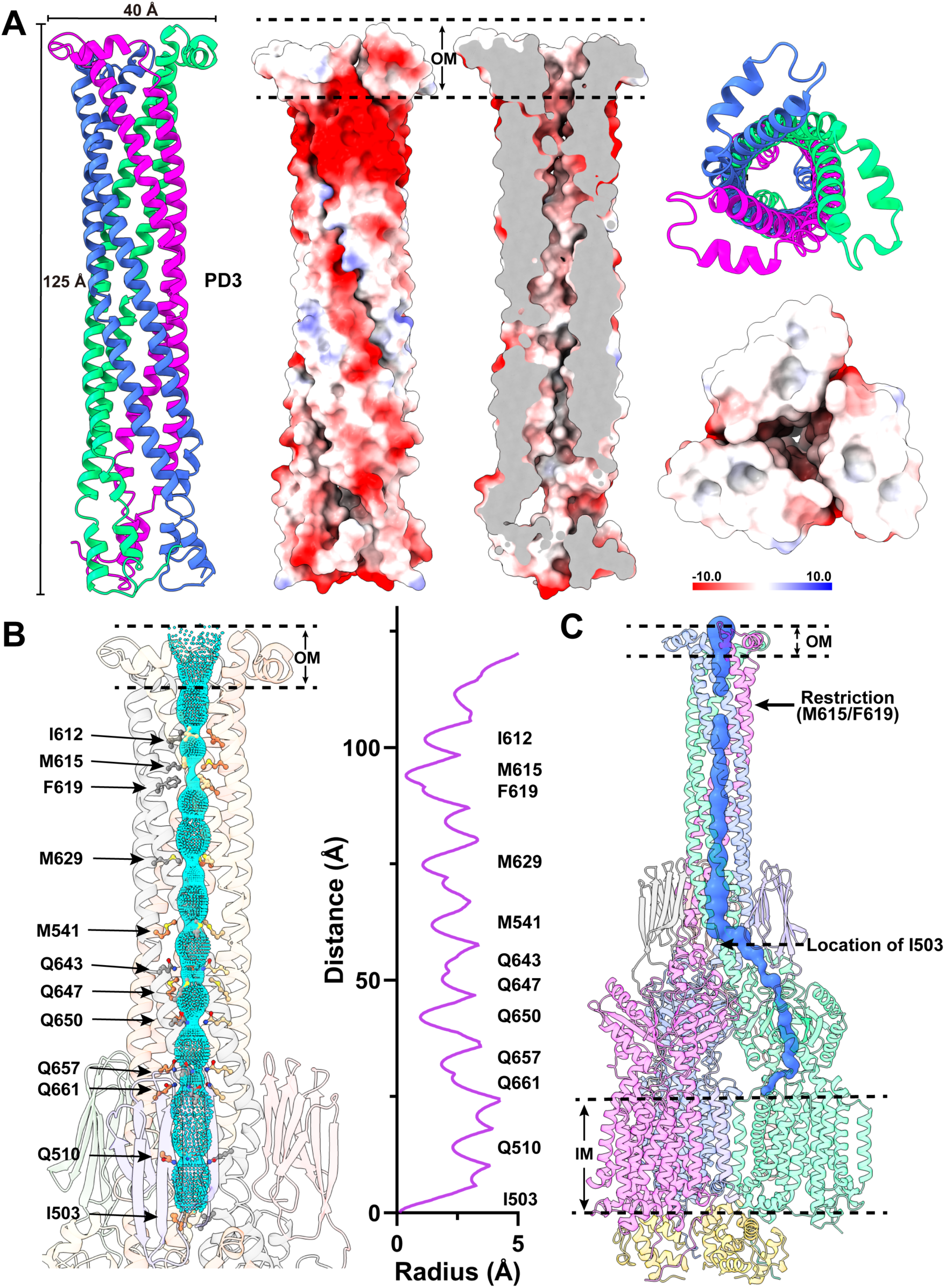
Structure of the trimeric PD3 channel. (A) Ribbon diagrams and electrostatic surface potentials of the side and top views of the trimer PD3 channel. The three PD3 domains of the MmpL5 trimer are in different colors. The blue (> 10 *k*B*T*) and red (< −10 *k*B*T*) colors of the electrostatic surface potential indicate the positively and negatively charged areas of the protein, respectively, where *k*B is the Boltzmann constant and *T* is absolute temperature. White denotes area between 10 *k*B*T* and −10 *k*B*T*. The needle-like apparatus of the PD3 channel is 125 Å long and 40 Å wide. (C) The interior of the central channel creates a pathway for transporting substrates. The channel, calculated by HOLE (http://www.holeprogram.org), is indicated by cyan dots. Residues forming the restriction sites and narrowest regions of the channel are represented with sticks. (D) The elongated substrate transport channel connecting the outer leaflet of the inner membrane up to the outer mycomembrane. The beginning of this channel is generated by a hydrophobic pocket created by TM7-TM10 of MmpL5. This channel (colored blue) was calculated using the program CAVER (http://loschmidt.chemi.muni.cz/caver). In both (B) and (C), the restriction site is created by residues M615 and F619 of each PD3 subdomain of the MmpL5 trimer.

Based on the structural information, this central channel is closed in conformation. The narrowest region of the channel, which forms a restriction site, is located at residues M615 and F619 of each MmpL5 protomer. At this site, the three sets of M615 and F619 residues of trimeric MmpL5 use mainly hydrophobic interactions to close this channel (Fig. 3B). The inner surface of the channel is predominantly neutral in charge, as indicated by the electrostatic surface potential (Fig. 3A), suggesting that the interior surface of the channel may be more favorable to bind and export hydrophobic substrates. Interestingly, the central channel created by the needle-like structure of subdomains PD3 of the MmpL5 trimer can be directly connected to the elongated channel spanning the pocket surrounded by TMs 7-10 at the outer leaflet of the cytoplasmic membrane and the cavity generated between subdomains PD1 and PD2 in the periplasmic domain of each MmpL5 protomer (Fig 3C). This observation indeed suggests that the assembled MmpL5 trimer by itself may be able to transport substrates across both the inner and outer membranes of the mycobacterium without requiring accessory proteins to facilitate the transport process.

#### Structure of the MmpS5 adaptor

We obtained the first structural information of the full-length MmpS5 adaptor within the AcpM-MmpL5-MmpS5 tripartite complex. The structure of MmpS5 presents as a single-spanning transmembrane protein, which contains an N-terminal transmembrane domain of one single TM helix and a C-terminal periplasmic domain of eight β strands. These two domains are connected by a flexible random loop of 20 amino acids. Thus, MmpS5 forms an elongated protein of 120 Å in length that spans the inner membrane and possesses an all β strand periplasmic domain (Fig 2D). Intriguingly, the structure of the tripartite complex depicts that the TM helix of one MmpS5 molecule solely interacts with TM8 of protomer 1 of MmpL5 within the trimer through hydrophobic interactions (Fig 4). However, the all β strand periplasmic domain of MmpS5 predominantly contacts protomers 2 and 3 of the MmpL5 trimer via hydrogen-bond, charge-charge, and dipole-dipole interactions (Fig 4). In addition, residues of the loop connecting the TM and β strand domains of MmpS5 are engaged in performing hydrogen-bond, charge-charge, dipole-dipole, and hydrophobic interactions with MmpL5 to further strengthen the binding (Fig 4). In this manner, each MmpS5 molecule appears to interlock and secure the trimeric oligomerization of the MmpL5 transporter. In addition, this structure demonstrates that the presence of the MmpS5 adaptor may be required for the establishment of the α-helical hairpin of each PD3 subdomain and the needle-like trimeric PD3 channel of the MmpL5 trimer.

**Fig 4.**
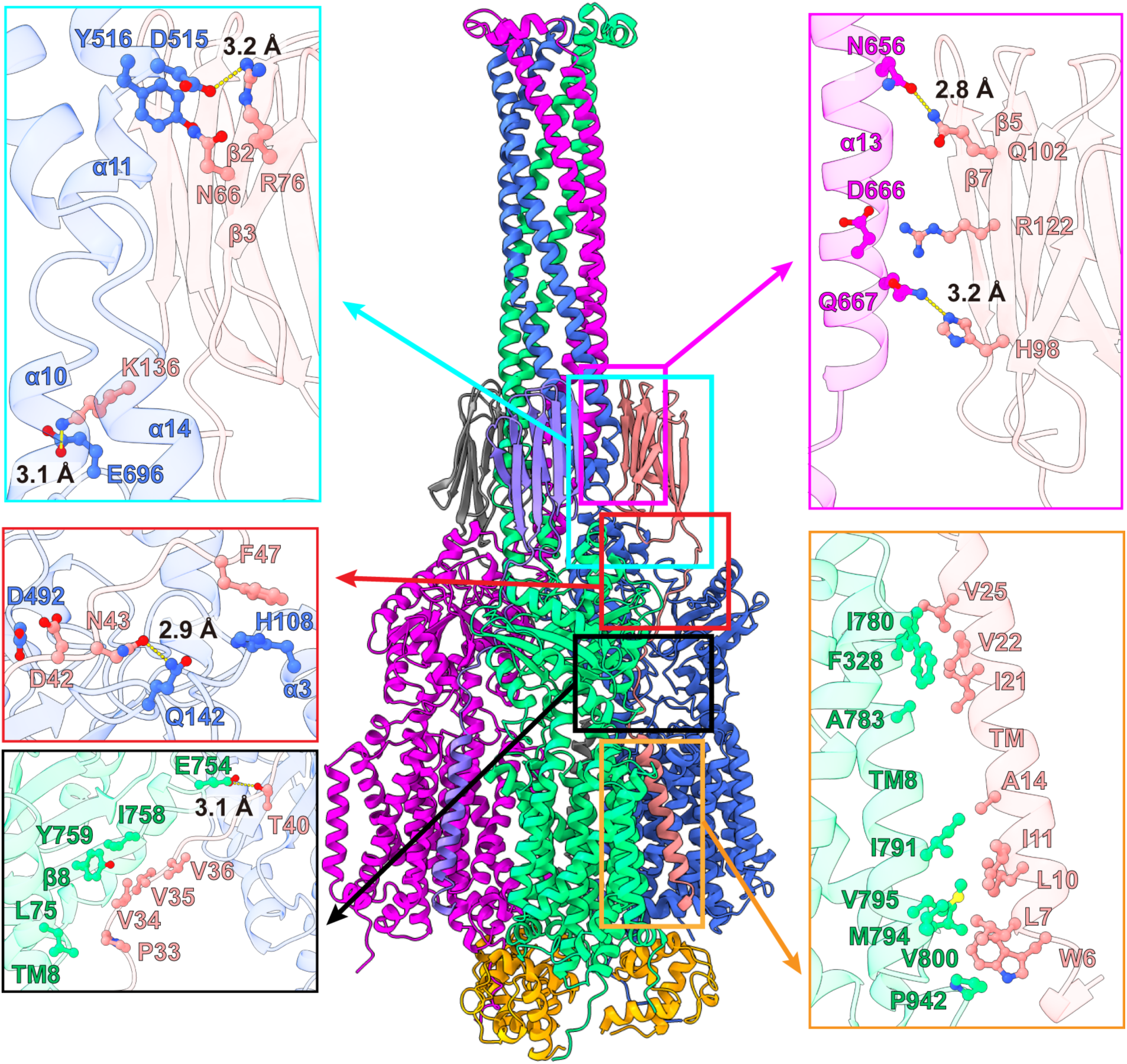
MmpL5-MmpS5 interactions. Protomers 1, 2 and 3 of MmpL5 within the trimer are colored green, blue and magenta, respectively. Molecules 1, 2 and 3 of MmpS5 are colored salmon, gray and slate, respectively. The three AcpM molecules are colored orange. The TM helix of molecule 1 of MmpS5 (colored salmon) solely interacts with TM8 of protomer 1 of MmpL5 through hydrophobic interactions. The all β strand periplasmic domain of molecule 1 of MmpS5 predominantly contacts protomer 2 of MmpL5 via hydrogen-bonds and dipole-dipole interactions. This all β strand domain of molecule 1 of MmpS5 also interacts with protomer 3 of MmpL5 via hydrogen-bonds and charge-charge interactions. In addition, residues of the loop region of molecule 1 of MmpS5 contact protomers 1 and 2 of MmpL5 through hydrogen-bonds, charge-charge, dipole-dipole, and hydrophobic interactions to further strengthen the binding. Residues participating in these interactions are in sticks. The hydrogen-bonds are indicated with dotted lines.

### The dipartite AcpM-MmpL5 trimeric complex

The third most highly populated particle images also belong to a membrane protein complex. We obtained 25,519 single-particle counts for this protein class in our cryo-EM data set. We identified this protein complex as the dipartite AcpM-MmpL5 complex, in which three of these AcpM-MmpL5 complex molecules assemble as a trimer. We then solved the cryo-EM structure of this membrane protein complex to a resolution of 3.37 Å (Fig 5A-C, Fig S1 and Table S1). The stoichiometry of this trimeric AcpM-MmpL5 complex is observed to have a 3:3 molar ratio.

**Fig 5.**
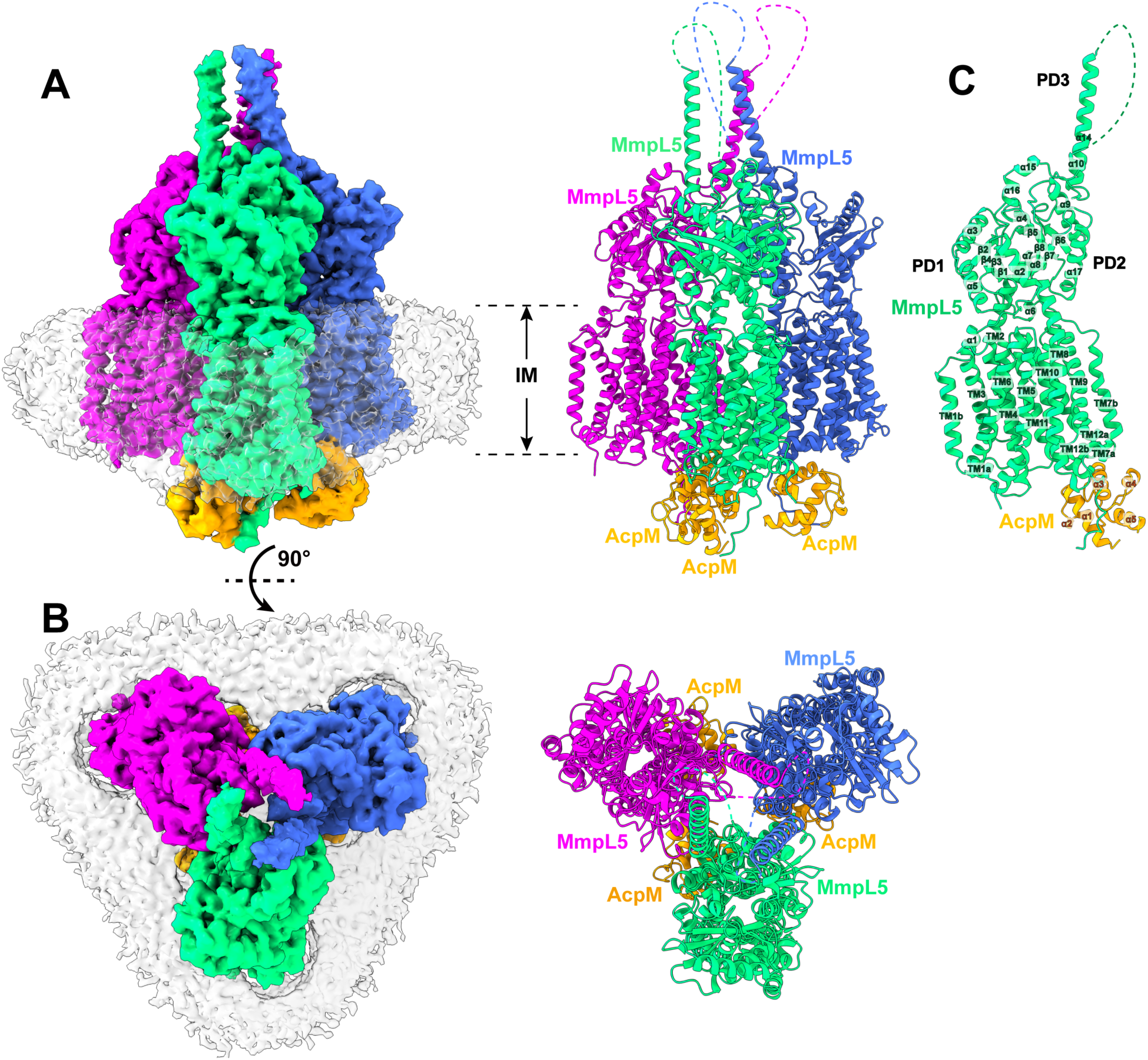
Structure of the dipartite AcpM-MmpL5 complex. (A) Side view and (B) top view of the cryo-EM maps and ribbon diagrams of the AcpM-MmpL5 complex. The structure indicates that MmpL5 assembles as a trimer. The stoichiometry of AcpM:MmpL5 within the complex is in the form of a 3:3 molar ratio. Individual MmpL5 molecules are colored differently. The AcpM molecules are colored orange and the detergent belts in the cryo-EM maps are colored gray. The inner membrane region is indicated as IM. (C) Structure of a single unit of AcpM-MmpL5 within the trimeric complex. The structure indicates that the top half of the PD3 hairpin is unstructured.

Similar to the structure of the AcpM-MmpL5-MmpS5 complex detailed above, MmpL5 exists as a homotrimer in this AcpM-MmpL5 complex (Fig 5A-B), where each MmpL5 protomer within the trimer is identical in conformation. In comparison with the AcpM-MmpL5-MmpS5 and AcpM-MmpL5 complexes, the major difference between these two MmpL5 structures arises from subdomain PD3 (Fig 5C). In the tripartite complex of AcpM-MmpL5-MmpS5, a full-length trimeric needle-like apparatus made up of the hairpin of subdomain PD3 from each MmpL5 protomer is established in the middle of the trimer. In the dipartite AcpM-MmpL5 complex, only the C-terminal half of α10 (residues 656-684) forming part of this trimeric needle can be easily seen, as the cryo-EM densities originating from residues 499-655 comprising the majority of this needle are difficult to trace from the cryo-EM map. It appears that the secondary structural elements of these residues may not be stable and are likely unstructured. The structural information suggests that the presence of MmpS5 may be a prerequisite for stabilizing the secondary α-helical structure of subdomain PD3 of MmpL5 with the MmpL5 trimer.

The AcpM-MmpL5 trimeric complex structure also shows that the interactions between AcpM and MmpL5 are almost the same as those seen in the AcpM-MmpL5 2D arrays. These interactions are mainly governed by hydrogen-bond, salt-bridge and electrostatic interactions.

## Discussion

The genome of Mtb encodes 13 MmpLs (*5*), which are critical for the mycobacterium’s physiology and pathogenesis. Several of these transporters, including MmpL3, MmpL7, MmpL8, MmpL10 and MmpL11, participate in the export of fatty acids and other lipid components, such as TMMs, MWEs, PMIDs, to strengthen the mycobacterial cell envelope. Additionally, some of these MmpLs have been proposed to form drug efflux pumps that mediate resistance to antituberculosis agents. For example, disrupting the MmpL5-MmpS5 system has been reported to result in the mycobacterium more susceptible to bedaquiline, clofazimine, rifabutin and azoles (*27, 29, 30*). The expression of MmpL5-MmpS5 is regulated by the transcriptional regulator Rv0678 (*31*), where its crystal structure indicates that it is a MarR-like regulatory protein. It has also been found that mutations in Rv0678 of Mtb is a major cause of resistance to bedaquiline (*32*). Furthermore, MmpL7 has been shown to contribute to isoniazid, rifampin and ethionamide resistance (*33*). In addition to cell wall biogenesis and drug resistance, MmpL5-MmpS5 forms a redundant system with MmpL4-MmpS4 to export siderophores, including mycobactins and carboxymycobactins, for iron acquisition (*15, 34*). Inactivation of both *mmps5* and *mmpL5* genes has been observed to reduce bacterial burden in the lungs of infected mice by more than 20,000-fold, resulting in the elimination of any visible lung pathology (*15*). Despite the importance of these MmpLs for mycobacterial physiology, virulence and pathogenesis, the mechanisms on how these transporters facilitate the transport of substrates, and even their functional oligomerizations and assemblies, remain unclear.

In this study, we overexpressed MmpL5 and MmpS5 in *M. smegmatis* to elucidate the structure and assembly of these two Mmp proteins from crude mycomembranes. From this membrane sample, we identified that MmpL5 specifically interacts with both AcpM and MmpS5 to form the AcpM-MmpL5-MmpS5 tripartite system. AcpM associates with MmpL5 in the cytoplasm, whereas MmpS5 contacts MmpL5 at both the transmembrane and periplasmic regions. This tripartite complex structure allowed us to visualize that MmpL5 is trimeric in form, something previously unknown and debated. The three MmpS5 adaptor molecules appear to be actively engaged in fastening the three MmpL5 protomers to secure the trimeric oligomerization. Within the MmpL5 trimer, subdomain PD3 of each MmpL5 protomer expands outward from subdomain PD2 to create a 125-Å long hairpin. These three hairpins interact with each other via coiled-coil interactions to further stabilize the trimeric organization of MmpL5. The three PD3 subdomains also create an elongated needle-like channel, which extends towards the outer mycomembrane and forms a passageway for substrate transport. The vertical dimension of the MmpL5 trimer is approximately 215 Å in length, which is probably long enough to span the entire mycobacterial cell envelope. Therefore, the MmpL5 trimer by itself may be capable of shuttling substrates across the mycobacterial membranes and it may not require additional accessory proteins to be involved in this transport process.

Our cryo-EM data indicate that MmpL5 can form two different oligomerization states and assemble into three distinct complexes with AcpM and MmpS5. The majority of the MmpL5 is associated with and creates two-dimensional protein arrays, where each protein unit within the arrays is identified as the AcpM-MmpL5 complex in the form of one molecule of AcpM and one molecule of MmpL5. In this arrangement, it is believed that the AcpM-MmpL5 units can freely diffuse laterally and move horizontally side-to-side across the inner membrane of the mycobacterium, where the 2D array assembly catalyzes and facilitates this diffusion process in parallel to the membrane surface. This observation is indeed in good agreement with single-molecule fluorescence measurement that MmpL5 can exist as a monomer, which showed substantial lateral displacements on the membrane plane (*23*). The second class of cryo-EM images reveals that each single particle is composed of an MmpL5 trimer, three AcpM and three MmpS5 molecules. This AcpM-MmpL5-MmpS5 complex is trimeric in oligomerization and assembles as a syringe and needle-like apparatus that extends from the inner membrane to the outer mycomembrane and spans the entire mycobacterial cell envelope. Presumably, this apparatus may be mainly stationary with very limited lateral movement, as both the outer and inner membranes may create this limitation. The third class of cryo-EM particles depicts a dipartite AcpM-MmpL5 complex, similar to that seen in the first class of cryo-EM images. However, in this third class, MmpL5 assembles as a trimer in the AcpM-MmpL5 complex. The assembly of the MmpL5 trimer in the AcpM-MmpL5 complex is similar to that found the AcpM-MmpL5-MmpS5 tripartite complex, but the secondary structural elements of the three PD3 periplasmic subdomains of the MmpL5 trimer are only form the bottom half of the elongated trimeric channel or the needle-like apparatus. The presence of MmpS5 may be a requirement to establish the full-length trimeric channel.

Based on these observations, we propose that AcpM-MmpL5 tends to form two-dimensional, monomeric arrays at the inner membrane of the mycobacterium in the absence of MmpS5. These 2D arrays may facilitate lateral diffusion of AcpM-MmpL5 protein molecules along the membrane plane, allowing them to effectively search for their partner MmpS5 adaptor proteins (Fig 6). Within the 2D arrays, the subdomain PD3 of each MmpL5 molecule is very flexible and forms a random loop between the N and C terminal halves of subdomain PD2. In the presence of MmpS5, MmpL5 can assemble into its trimeric oligomerization, where three MmpS5 adaptor molecules actively participate in anchoring three MmpL5 transporter molecules together to form the AcpM-MmpL5-MmpS5 complex in a molar ratio of 3:3:3 (Fig 6). This trimeric oligomerization also stabilizes the secondary structure of subdomain PD3 of each MmpL5 molecule to form an all α-helical hairpin and protrude towards the outer mycomembrane of the mycobacterium. These three hairpins further strengthen the trimeric oligomerization by forming an elongated channel via coiled-coil interaction, where the trimeric channel also creates a passageway for transporting substrates. Once fully assembled, the vertical dimension of trimeric MmpL5 is long enough to span the entire periplasmic space so that the tip of this elongated channel can insert into the outer mycomembrane. The elongated, syringe and needle-like structure of the MmpL5 trimer also suggests that this membrane protein is capable of spanning both the inner and outer membranes of the mycobacterium to transport substrates across the entire cell envelope. As the mycobacterial cell envelope contains the arabinogalactan and peptidoglycan layers, significant intrinsic resistance likely prevents the lateral movement of the trimeric AcpM-MmpL5-MmpS5 complex on the membrane plane. Therefore, this trimeric AcpM-MmpL5-MmpS5 complex should be more or less stationary in position. This finding is indeed in good agreement with results from fluorescence experiments that show that the trimeric MmpL5-MmpS5 complex is restricted with limited horizontal diffusion (*23*). After the extrusion of substrates, the trimeric AcpM-MmpL5-MmpS5 complex can be dismantled. The first step of the disassembly process may be the removal of the MmpS5 adaptor molecules to form the trimeric AcpM-MmpL5 complex (Fig 6). Based on the single-particle counts, this state is much less populated in comparison with the trimeric AcpM-MmpL5-MmpS5 form. Therefore, the trimeric AcpM-MmpL5 complex may represent a less stable intermediate state. At this state, the secondary structural elements of subdomain PD3 of MmpL5 become unsteady and the top half of the secondary structure of the hairpin becomes untraceable. The next step is that the trimeric AcpM-MmpL5 complex continues to dismantle to form three monomeric AcpM-MmpL5 complexes. At this stage, the secondary structural elements of the entire PD3 subdomain are untraceable and form a flexible loop. These AcpM-MmpL5 monomers may then gather to reform 2D arrays to stabilize the AcpM-MmpL5 complex structure (Fig 6).

**Fig 6.**
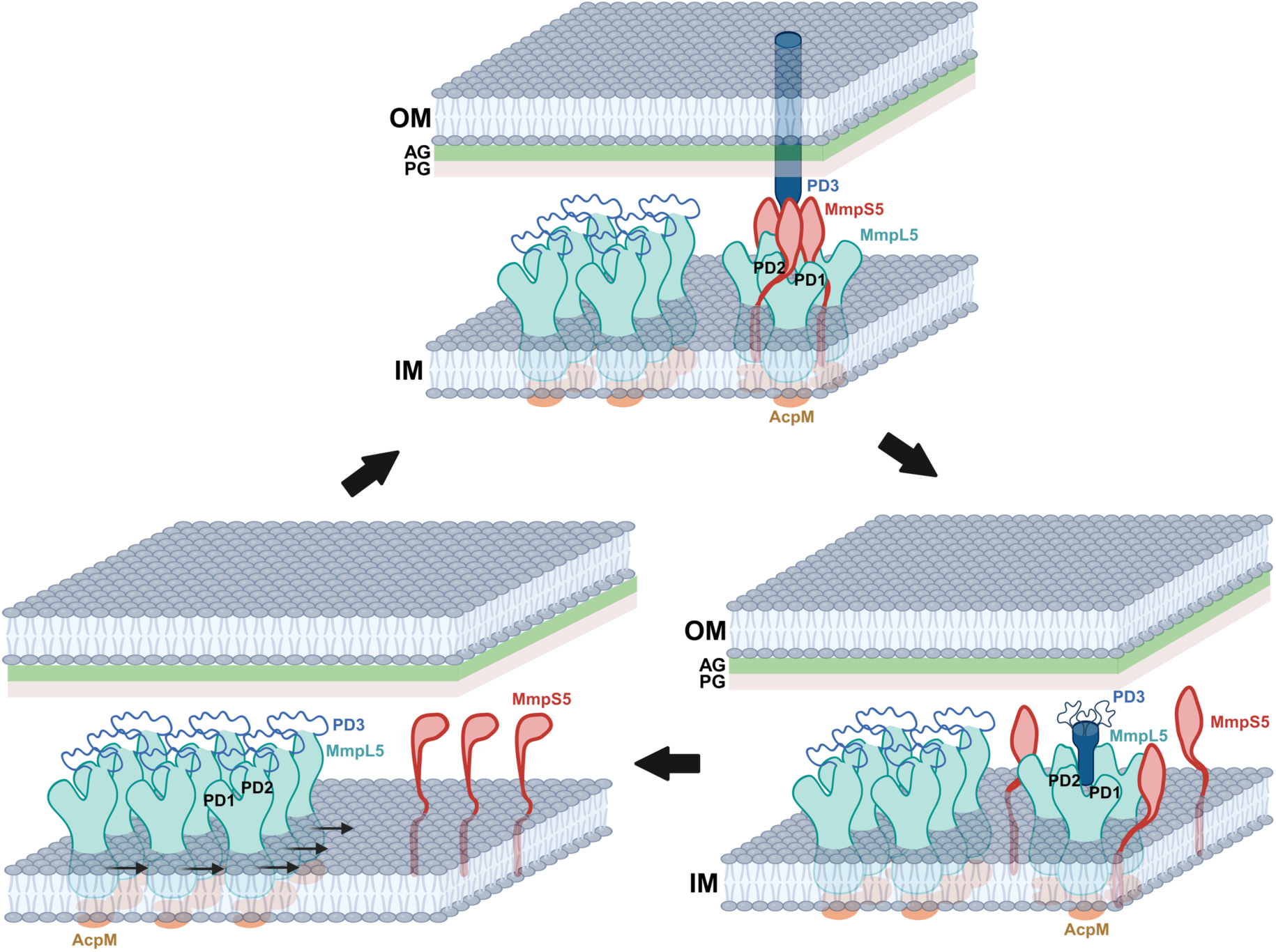
Proposed mechanism for the assembly of the AcpM-MmpL5-MmpS5 trimeric complex. In the absence of MmpS5, the AcpM-MmpL5 molecules form 2D arrays, which facilitate their lateral diffusion along the inner membrane of the mycobacterium. In the presence of MmpS5, MmpL5 forms a trimer and the tripartite AcpM-MmpL5-MmpS5 complex assembles. The trimer oligomerization of MmpL5 and the interaction between MmpL5 and MmpS5 stabilize the secondary structural elements of the three PD3 subdomains of the MmpL5 trimer, allowing them to create an elongated channel which protrudes towards the outer mycomembrane. The presence of the arabinogalactan (AG) and peptidoglycan (PG) layers create sufficient resistance to prevent the lateral movement of the trimeric AcpM-MmpL5-MmpS5 complex on the membrane plane. After the extrusion of substrates, the trimeric AcpM-MmpL5-MmpS5 complex can be dismantled by removing the three MmpS5 adaptor molecules to form the trimeric AcpM-MmpL5 complex. In this state, the secondary structural elements of subdomain PD3 of MmpL5 become unsteady and the top portion of the secondary structure of the hairpin becomes untraceable. The trimeric AcpM-MmpL5 complex then dismantles to form three monomeric AcpM-MmpL5 complexes, where the secondary structural elements of the entire PD3 subdomain of each MmpL5 molecule are untraceable and form a flexible loop. These AcpM-MmpL5 monomers then repack and form 2D arrays to stabilize the AcpM-MmpL5 complex structure.

However, the AcpM-MmpL5 trimer may be present before the complete formation of the AcpM-MmpL5-MmpS5 complex. In this case, the AcpM-MmpL5 trimeric complex would be the intermediate state between the dipartite AcpM-MmpL5 monomeric and tripartite AcpM-MmpL5-MmpS5 trimeric states. Nonetheless, our cryo-EM structures allow us to observe the trimeric oligomerization of the MmpL5 transporter, where the presence of the MmpS5 adaptor facilitates the MmpL5 timer to form a machinery that is able to span the entire mycobacterial cell envelope. The structural information of the AcpM-MmpL5-MmpS5 complex detailed here likely has a much wider implication beyond just the MmpL family of transporters. For instance, the most recent reported structure of PfNCR1 (*35*), an RND-type cholesterol transporter that regulates membrane homeostasis of the malarial parasite *Plasmodium falciparum*, indicates that this membrane protein is monomeric in oligomerization. There is a good chance that this transporter may assemble into a higher ordered oligomerization state to function when it contacts its accessory proteins and/or interaction partners.

## Methods

### Protein expression and membrane preparation

The *M. smegmatis* MmpS5 (MSMEG_1381) and MmpL5 (MSMEG_1382) genes were cloned into the pBUN250 expression vector in frame with a 6xHis tag at the C-terminus of MmpL5. The MmpS5 and tagged MmpL5 proteins were overproduced in *M. smegmatis* MC^2^-155. Cells were grown in 12 L of LB medium supplemented with 0.5% glycerol, 0.05% Tween-80, and 50 μg/ml kanamycin at 37°C overnight. The culture was then treated with 0.2% acetamide and grown at 18°C during induced expression. Cells were harvested after 20 h of induction. The collected bacteria were resuspended in low salt buffer (100 mM sodium phosphate (pH 7.2), 10% glycerol, 1 mM ethylenediaminetetraacetic acid (EDTA) and 1 mM phenylmethanesulfonyl fluoride (PMSF)) and disrupted with a French pressure cell. The membrane fraction was collected, washed twice with low salt buffer and twice with final buffer (20 mM HEPES-NaOH buffer (pH 7.5) and 20 mM NaCl). The membrane was then solubilized in 2% (w/v) lauryl maltose neopentyl glycol (LMNG) overnight at 4°C. Insoluble material was removed by ultracentrifugation at 186,000 x g.

### Membrane protein enrichment

The membrane protein component of the solubilized membrane was enriched using sucrose cushion ultracentrifugation. Briefly, a 2 ml sucrose buffer (45% sucrose, 20 mM Tris-HCl (pH 7.5), 100 mM NaCl, 0.0001% LMNG) was added to the bottom of the centrifugation tube. A 2 ml detergent solubilized sample was slowly added to the top of the sucrose buffer and centrifuged at 100,000 x g for 16 h at 4 °C. After centrifugation, the membrane protein pellet was dissolved in a buffer containing 20 mM Tris-HCl (pH 7.5), 20 mM NaCl and 0.002% LMNG for cryo-EM sample preparation.

### Cryo-EM sample preparation

2.5 μl of solubilized sample (8 mg/ml) from the sucrose cushion was directly applied to glow-discharged holey carbon grids (Quantifoil Cu R1.2/1.3, 300 mesh), blotted for 12 s and then plunge-frozen in liquid ethane using a Vitrobot (Thermo Fisher). The resulting grids were then transferred into cartridges prior to data collection.

### Data Collection

Samples were collected at the Cleveland Center for Membrane & Structural Biology (CCMSB) using a Titan Krios cryo-electron transmission microscope (Thermo Fisher Scientific). Images were recorded using 0.8 −1.5 μm defocus on a K3 direct electron detector (Gatan) using super resolution at 81,000 x magnification. The super resolution size was 0.535 Å/pixel corresponding to a physical pixel size of 1.07 Å/pixel (Table S1).

### Data Processing

Initial image stacks were binned by a factor of 2 and motion corrected using cryoSPARC v3 (*36*). The patchCTF function was used to estimate the contrast transfer function (*36*). The Topaz tool (*37*), default ResNet16 (64 U) pre-trained model, was used to pick initial particle sets. The BaR protocol (*28*) was used to separate different initial structure classes. First, during the “build” phase, particles are sorted via multiple rounds of 2D classification to identify particles with distinct structural features. Particle stacks consisting of distinct 2D representations are refined and grouped together to form starting 3D *ab initio* models. All starting models are built using C1 symmetry to reduce bias. Starting models that contain structurally viable global features are then used as filters in a series of heterogeneous refinements using the original particle stack. At the “retrieve” phase, each model was retrieved additional particles relative to the initial stack, including a more diverse set of orientations. These refined particle stacks are then used to generate refined 3D models that can be used as updated filters to retrieve even more particles belonging to each distinct class. Once the build and retrieve iterations no longer add additional particles, non-uniform refinement is used on the final 3D model to generate a high-resolution density map. Protein identification of the protein complexes consisting of AcpM and MmpL5 with or without MmpS5 were done by a combination of manual building and utilizing the program DeepTracer (*38*) to obtain amino acid sequences of proteins that assemble with each other to form these protein complexes. The presence of the MmpL5, MmpS5 and AcpM proteins in our sample was also confirmed by proteomics (Table S2).

### Model building and refinement

Model building of all protein complexes were performed using Coot (*39*) based on their respective cryo-EM maps. Structural refinements were accomplished using the phenix.real_space_refine program (*40*) from the PHENIX suite (*41*). The final atomic models of these protein complexes were evaluated using MolProbity (*42*). The statistics associated with data collection, 3D reconstruction and model refinement are included in Table S1.

### Accession codes

Atomic coordinates and EM maps for monomeric AcpM-MmpL5, trimeric AcpM-MmpL5-MmpS5 and trimeric AcpM-MmpL5 have been deposited with PDB accession codes 9MX0, 9MVZ and 9MW0, and EMDB accession codes EMD-48706, EMD-48675 and EMD-48676.

## Acknowledgements

This work was supported by an NIH Grant R01AI187294 (E.W.Y.).

