## Supplementary material for "Structures of MmpL complexes reveal the assembly and mechanism of this family of transporters": Fig S1 and Tables S1-S2

**Supplementary Information**

**Secondary structural elements of MmpL5**

The TMs, α-helices and β-strands of MmpL5 are assigned numerically from the N- to C-termini: TM1 (25-51)), α1 (51-62), α2 (72-84), β1 (91-99), α3 (105-120), β2 (126-128), α4 (138-141), β3 (148-155), α5 (162-176), β4 (184-189), TM2 (191-225), TM3 (228-255), TM4 (262-293), TM5 (297-328), TM6 (332-365), TM7 (a (379-390) and b (392-412)), α6 (420-422), α7 (428-439), α8 (442-445), β5 (448-453), α9 (461-476), β6 (481-484), α10 (499-505), α11 (508-575), α12 (581-589), α13 (599-672), α14 (677-684), α15 (695-699), α16 (701-710), β7 (717-723), α17 (731-748), β8 (757-761), TM8 (763-797), TM9 (800-826), TM10 (837-866), TM11 (868-899) and TM12 (903-935).

**Secondary structural elements of MmpS5**

The TM and β-strands of MmpLS are assigned numerically from the N- to C-termini: TM (6-27), β1 (50-58), β2 (63-68), β3 (74-78), β4 (84-90), β5 (98-103), β6 (107-114), β7 (117-127) and β8 (130-134).

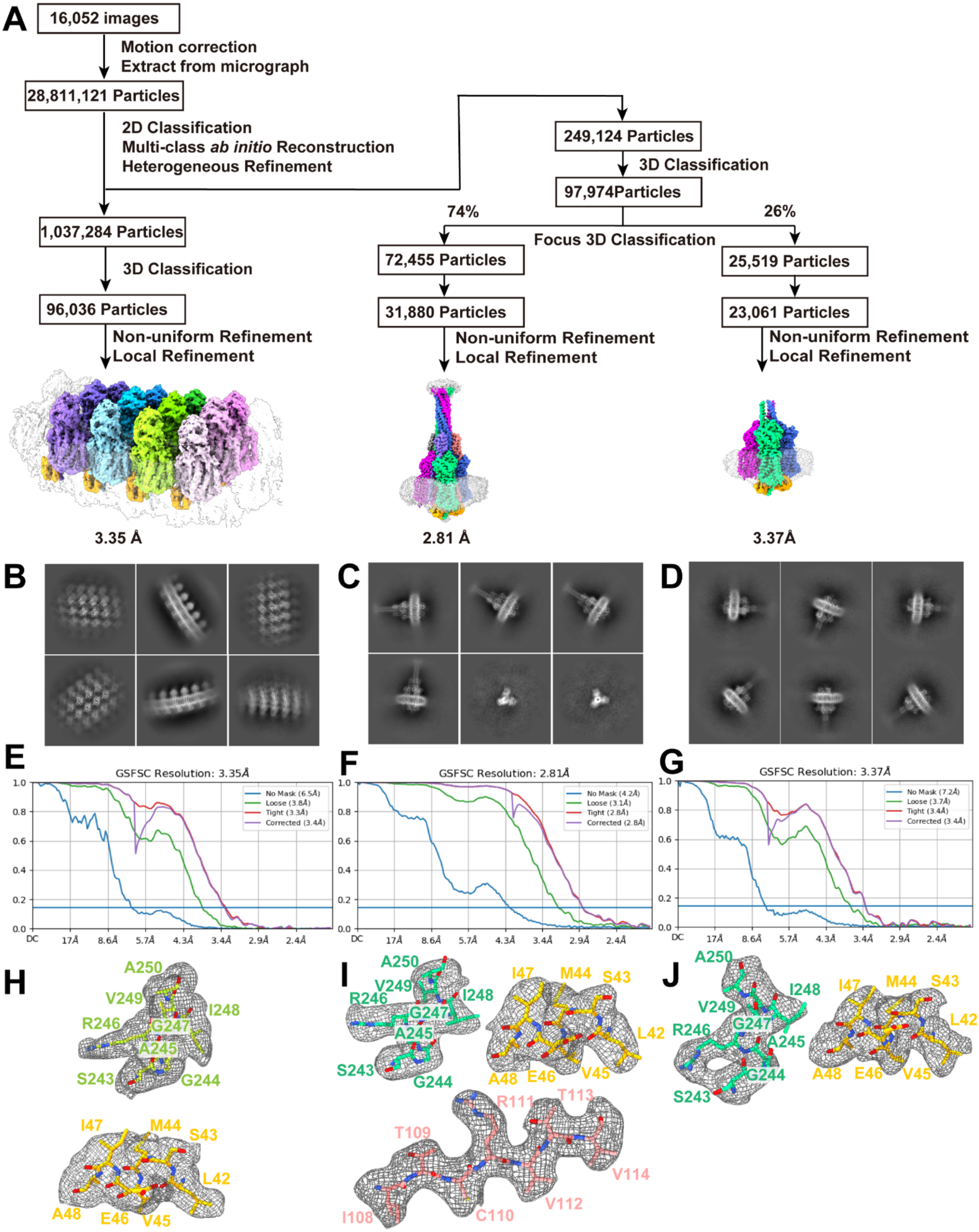


Fig S1. **Build and Retrieve workflow.** Cryo-EM processing begins as standard workflow, where motion-corrected micrographs are picked, particles undergo 2D classification and initial models are iteratively built. These low-resolution initial models are then used to retrieve particles from the cleaned dataset, resulting in 3 high-resolution maps. These maps lead us to solve cryo-EM structures of the AcpM-MmpL5 complex in 2D arrays, the AcpM-MmpL5-MmpS5 complex and the trimeric AcpM-MmpL5 complex to resolutions of 3.35 Å, 2.81 Å and 3.37 Å, respectively.

**Table S1. Cryo-EM data collection and refinement statistics.**

| **Data collection** | **Sucrose cushion sample of *M. Smegmatis* membrane expressed with MmpL5/S5** | | |
|---|---|---|---|
| Magnification | | 81,000 | |
| Voltage (kV) | | 300 | |
| Electron Microscope | | Krios-GIF-K3 | |
| Defocus (um) | | -0.8 to -1.5 | |
| Energy filter width (eV) | | 20 | |
| Pixel size (Å) | | 1.07 (0.535) | |
| Total dose ($e^-$/ $Å^2$) | 40.2 | 50 | 50 |
| Number of frames | 46 | 40 | 40 |
| Number of micrographs | 3,855 | 6,201 | 5,996 |
| Number of Initial particles | | 28,811,121 | |
| **Refinement** | **Monomeric AcpM-MmpL5** | **Trimeric AcpM-MmpL5-MmpS5** | **Trimeric AcpM-MmpL5-MmpS5** |
| Number of total particles | 96,036 | 31,880 | 23,061 |
| GS-FSC Resolution (0.143, Å)[a] | 3.35 | 2.81 | 3.37 |
| <u>Model composition</u> | | | |
| Chains | 32 | 9 | 6 |
| Protein residues | 9,684 | 3,468 | 2,589 |
| Ligand | 8 | 3 | 3 |
| <u>r.m.s.d.</u> | | | |
| Bond lengths (Å) | 0.002 | 0.005 | 0.003 |
| Bond angles (°) | 0.388 | 0.520 | 0.475 |
| **Validation** | | | |
| MolProbity score | 1.62 | 1.51 | 1.48 |
| Clash score | 7.42 | 4.43 | 5.27 |
| <u>Ramachandran plot</u> | | | |
| Favored (%) | 98.94 | 98.41 | 98.87 |
| Allowed (%) | 1.06 | 1.59 | 1.13 |
| Disallowed (%) | 0.00 | 0.00 | 0.00 |
| CC Mask | 0.66 | 0.81 | 0.81 |

[a]Gold-Standard Fourier-Shell Correlation

**Table S2 Proteomic analysis.**

| Ranking | Protein ID | Accession | Mass kDa | Peptide unique | Seq. Coverage (%) | Sequest Score |
|---|---|---|---|---|---|---|
| 1 | **MmpL5 protein** | **A0QS80** | **105.4** | **60** | **54.0** | **4290.09** |
| 2 | Cytochrome bc1 complex Rieske iron-sulfur subunit | I7GD61 | 46.3 | 35 | 66.0 | 1328.80 |
| 3 | ATP-dependent zinc metalloprotease FtsH | I7GG40 | 83.5 | 39 | 62.0 | 1060.13 |
| 4 | cytochrome-c oxidase | I7GD63 | 38.0 | 28 | 58.0 | 925.52 |
| 5 | Elongation factor Tu | A0QS98 | 43.7 | 22 | 78.0 | 874.20 |
| 6 | NADH:ubiquinone reductase (non-electrogenic) | I7GBR7 | 49.0 | 22 | 71.0 | 840.85 |
| 7 | ATP synthase subunit beta | A0R200 | 51.6 | 30 | 88.0 | 834.46 |
| 8 | UPF0182 protein MSMEG_1959/MSMEI_1915 | A0QTT7 | 109.0 | 36 | 53.0 | 820.69 |
| 9 | DNA-directed RNA polymerase subunit beta' | A0QS66 | 146.4 | 68 | 67.0 | 795.19 |
| 10 | Phage shock protein A, PspA | I7G0I2 | 30.3 | 22 | 90.0 | 714.75 |
| : | | | | | | |
| 14 | **MmpS5 protein** | **A0QS79** | **14.9** | **8** | **80.0** | **614.55** |
| : | | | | | | |
| 280 | **Meromycolate extension acyl carrier protein (AcpM)** | **A0R0B3** | **10.7** | **5** | **36.0** | **94.48** |
